# Exploring the Repetitive DNA Diversity in *Solanum betaceum* (Solanaceae)

**DOI:** 10.1101/2025.08.26.672483

**Authors:** MA Sader, M Vaio, A Trenchi, MM Jaramillo Zapata, F Chiarini, A López, JD Urdampilleta

## Abstract

The Solanaceae family, known for its diverse and economically important crops, includes the genus *Solanum*, which comprises 1,245 species. *Solanum betaceum* (tree tomato), native to the Andes and cultivated globally, is a promising species due to its nutritional value and market potential. The Cyphomandra clade, which includes the tree tomato, is characterized by huge genomes and chromosomes, with repetitive DNA elements (e.g., retrotransposons and satellite DNA) playing crucial roles in genomic and evolutionary studies. Despite its importance, genetic research on *S. betaceum* remains limited. This study addresses this knowledge gap by characterizing the repetitive DNA fraction to better understand intraspecific variation and develop molecular markers. Samples from five populations in northwestern Argentina were cultivated, and genome size was assessed via flow cytometry. Illumina HiSeq sequencing combined with RepeatExplorer analysis was used to identify repetitive DNA elements. Cytogenetic techniques, including CMA/DAPI staining and fluorescence in situ hybridization (FISH), were employed to detect satellite DNA patterns. Genome size analysis revealed slight variation among populations. Repetitive DNA accounted for 63.5% of the genome, with Ty3-gypsy retrotransposons being the most abundant (51.44%). Satellite DNA and rDNA were less prevalent, comprising 0.93% and 0.30% of the genome, respectively. Population comparisons showed consistent proportions of repetitive DNA overall, with notable differences in Ty3-gypsy-Tekay and satellite DNA fractions. This study provides a detailed profile of the repetitive DNA landscape in *S. betaceum*, uncovering intraspecific differences and delivering valuable genomic insights for future breeding and conservation efforts.

## Introduction

Solanaceae is a remarkable family of flowering plants, highly regarded by botanists and society. This diverse family encompasses essential crops alongside ornamental plants, weeds, and various species used as biological models. Within the Solanaceae family, the *Solanum* L. genus is one of the world’s largest and economically most important plant genera, with 1,245 currently accepted species. Twenty-four of these are major crops (Knapp et al., 2004), including the potato (*S. tuberosum* L.), tomato (*S. lycopersicum* L.), and brinjal/eggplant (*S. melongena* L.), as well as some lesser-known cultivated species, such as pepino (*S. muricatum* Aiton), lulo/naranjilla *(S. quitoense* Lam. and relatives), cocona (*S. sessiliflorum* Dunal), and tree tomato or tamarillo (*S. betaceum* Cav.) (Hilgenhof et al., 2023).

*Solanum betaceum*, (syn. *Cyphomandra betacea* (Cav.) commonly known as tree tomato, belongs to the Cyphomandra clade of *Solanum*, and is a fast-growing fruit-bearing plant native to the Andean regions of South America (Prohens & Nuez, 2001). It is a domesticated species grown between 600 and 3,200 meters above sea level (Bohs, 2015). The exact region of origin of *S. betaceum* is not known, but some wild or naturalized populations have been reported in southern Bolivia and northeastern Argentina and may be an indication of the area (Bohs, 1991; Brücher, 1977). The cultivation of tree tomatoes has expanded to various tropicals and subtropicals countries, including New Zealand, Australia, and India (Ramírez and Kallarackal, 2019). In recent years, interest in tree tomatoes has increased due to their abundant content of sugars, organic acids, minerals, vitamins (particularly vitamin C and B6), carotenoids, anthocyanins, and phenolics (Acosta-Quezada et al., 2015; Espin et al., 2016). The fruit is characterized by a slightly bitter, sour, and astringent taste accompanied by a distinctive aroma. Consequently, tree tomato has transitioned from a neglected crop to an up-and-coming fruit crop (Pacheco et al., 2021).

In Solanaceae, average chromosome size varies from approximately 1 μm (in *Solanum* and *Atropa*) to 6.5–11.5 μm (in *Cestrum* and the *Cyphomandra* clade of *Solanum*). However, most species have small to medium-sized chromosomes, with an overall mean chromosome length of 2.95 ± 1.78 μm (Deanna et al., 2020). *Solanum* is characterized by relatively small to medium-sized chromosomes compared to other Solanaceae and angiosperms (Badr et al., 1997; Guerra, 2000; Chiarini et al.,2018), with most species falling within the 1–3 μm range. The genome size of species within the *Cyphomandra* clade is notably large, with and average 1C = 10.80 pg compared to the average genome size in the *Solanum* genus 1C = 1.40 pg. The smallest genome size reported in the clade is that of *S. corymbiflorum* (Sendtn.) Bohs (1C = 6.80 pg), while the largest corresponds to *S. hartwegii* (1C = 24.80 pg) (Pringle and Murray, 1991; https://cvalues.science.kew.org/). These large genome sizes are directly correlated with larger chromosome sizes. Both large chromosomes and genome sizes represent exceptional synapomorphies of the *Cyphomandra* clade (Bohs, 1994, 2001; Chiarini, et al., 2017) supporting the value of these karyotype attributes for clade characterization. The number of CMA/DAPI bands and rDNA sites does not account for the observed differences in genome size. Therefore, the accumulation and dispersion of other repetitive sequences, such as transposable elements, may be associated with karyotype changes (Mesquita et al., 2024). These large synapomorphic chromosomes represent an interesting case for studying phenomena such as “genome obesity” (Bennetzen and Kellogg, 1997), characterized by increases in DNA content due to the incorporation of heterochromatin, repetitive sequences, and retrotransposons (Fregonezi et al., 2007; Miguel et al., 2012; Scaldaferro et al., 2013).

Repetitive DNA (repeats) forms the structural backbone of the chromosomes and represents the fastest-evolving sequences in the genome. Their rapid sequence turnover, along with genome-wide amplification and loss, can drastically impact karyotypes and overall genome composition. The resulting genetic diversity may facilitate species adaptation to environmental changes, potentially leading to speciation (Paço et al., 2015). Repetitive DNA also, constitutes a significant portion of plant genomes, accounting for approximately 50–90% of their total genomic content (Bennett et al., 1982; Kubis et al., 1998; Heslop-Harrison, 2000). This DNA comprises dispersed elements, such as transposable elements, and tandem sequences, like satellite DNA, both vary widely in abundance. Retrotransposons are a class of transposable element that propagate within a genome via an RNA intermediate. They can be classified into two major types: long terminal repeat (LTR) retrotransposons and non-LTR retrotransposons. LTR retrotransposons resemble retroviruses and include sequences that facilitate their integration into the genome. Tandem repeats, such as satellite DNAs (satDNAs) and ribosomal DNAs (rDNAs), are often located at key chromosomal regions including the (peri)centromeres and (sub)telomeres, as well as specific loci such as the ribosomal RNA genes, which often correspond to secondary constrictions when active, and large chromatin domains like constitutive heterochromatin (Biscotti et al., 2015a, b). Satellite DNA (satDNA), known for forming extensive arrays on chromosomes, often associated with heterochromatin regions, is detectable via fluorescence *in situ* hybridization (FISH) (Schwarzacher and Heslop-Harrison, 2000), facilitating cytogenetic studies. Key features of satDNA include monomer sizes typically larger than 100–150 bp and tandem arrays that can span up to 100 Mb (Sharma and Raina, 2005; Hemleben et al., 2007; Mehrotra and Goyal, 2014). Despite being regarded as non-coding sequences, their monomeric lengths often fluctuate between 150–180 and 320–360 bp, resembling structural motifs of mono- and dinucleosomes (Kubis et al., 1998; Macas et al., 2002). Advances in genomic sequencing techniques, such as next generation sequencing (NGS), combined with progress in bioinformatics, continue to reveal new insights into the structural diversity of satDNA (Ruiz-Ruano et al., 2016; Garrido-Ramos, 2017; Lower et al., 2018). Nonetheless, many fundamental aspects of satDNA biology remain enigmatic. The evolution of satDNA within genomes entails divergence in both sequence and copy number. The abundance of satDNA in eukaryotic genomes can fluctuate dramatically and rapidly across generations, resulting in considerable polymorphism in satellite array lengths (Plohl et al., 2012). Within an array, sequence identity evolves via concerted evolution, leading to monomers becoming more similar than expected due to stochastic changes (Dover, 1986; Plohl, 2010). Homogenization of monomers occurs more efficiently in sexually reproducing species, facilitated by meiotic recombination (Mantovani et al., 1997). Chromosome organization profoundly influences key cellular processes such as chromosome pairing, segregation, gene arrangement, and expression. In this context, repetitive DNA, including satDNA, plays crucial roles in DNA packaging and chromatin condensation (Heslop-Harrison, 2000). Given that satDNA distribution can facilitate in recognizing homologous chromosome pairs, alterations between lineages have often preceded speciation events (Hemleben et al., 2007). The similarities and disparities in genomic satDNA composition among species can be explain by the “library model,” which posits varied amplification of satDNA across independent lineages (Garrido-Ramos, 2015). Nonetheless, the sporadic distribution of certain satDNA types across eukaryotes (both animals and plants) suggests the possibility of multiple horizontal transfer events during evolution (Yang et al., 2020).

SatDNA commonly shows species-specific sequence types, abundance, and chromosome distribution patterns; however, its intraspecific characteristics have been poorly described. Despite tree tomatoes’ great potential as a major emerging fruit crop, no genetic or genomic studies have been conducted on this species to date. Therefore, the main objectives of this work were to characterize and describe the repetitive fraction of the tree tomato, identify both qualitative and quantitative aspects of intraspecific variation across different accessions, and contribute to the development of molecular markers. To this end, repeatome analyses were performed using low-coverage genome sequencing data to estimate the repeat fractions and to harness genomic differences among genotypes. Finally, by analyzing tandemly repeated sequences, we developed cytogenetic reference probes to characterize the karyotype.

## Materials and methods

### Plant material

Samples from five populations of *Solanum betaceum* were collected in northwestern Argentina (Fig1) and maintained in pots under controlled conditions in a growth chamber at 25 °C. Vouchers specimens were deposited in the herbarium of the Botanical Museum of Córdoba, Argentina (CORD).

**Figure 1.**
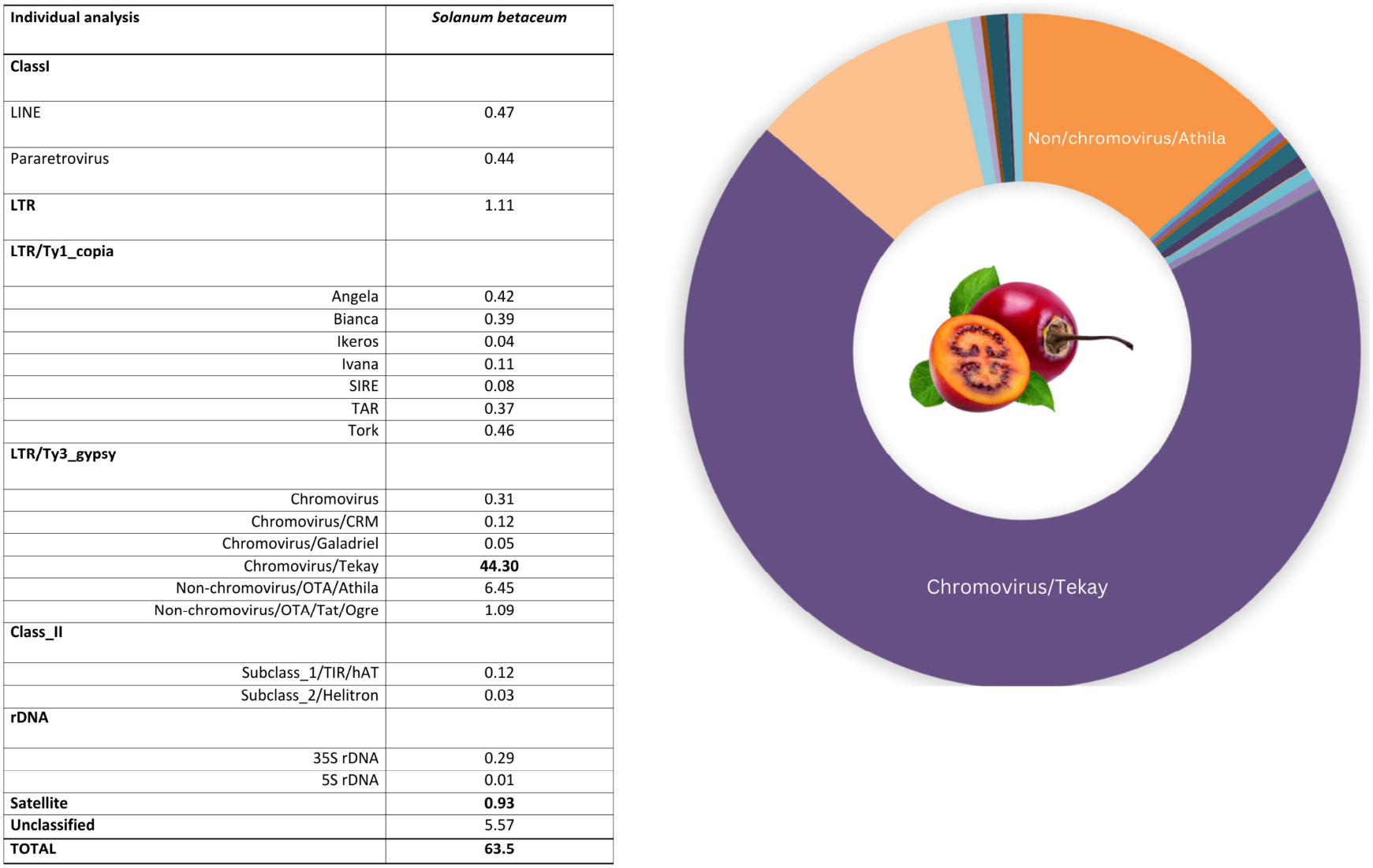
Representatives of distinct *Solanum betaceum* populations along with their corresponding voucher specimens

### DNA content estimations by flow cytometry

DNA measurements and nuclei extractions were performed following the procedure described by Galbraith et al. (1983) using the Woody Plant Buffer, WPB (Loureiro et al., 2007). Briefly, small pieces of leaves from the sample and the appropriate standard were mixed in a glass Petri dish (96 mm) with 1 mL WPB, co-chopped with a sharp razor blade, filtered through a 30 um mesh nylon membrane into a cytometry tube. Nuclei were stained with propidium iodide (50 µg/mL final concentration) and analyzed using a Partec CyFlow® Space flow cytometer (Sysmex-Partec GmbH, Münster, Germany) with data processing in the FloMax® program (Version 2.4d; Sysmex-Partec GmbH, Münster, Germany). *Pisum sativum* (1C = 4.90 pg; Dolezel et al. 1992), was used as an internal reference standard. At least three DNA estimations were carried out for each sample. Nuclear DNA content (2C) was calculated as: (Sample peak mean/Standard peak mean) x 2C DNA content of standard (pg).

### High□throughput sequencing, data processing, and repetitive DNA identification

Genomic DNA was isolated from silica gel-dried leaves following the CTAB II protocol (Weising et al. 2005). DNA integrity was assessed by electrophoresis on an agarose gel 1%. Low coverage sequencing (1x) was performed on an Illumina HiSeq using 150 bp paired-end reads by BGI Group (Hong Kong, China). Publicly available sequencing data for *S. betaceum* available online at the European Nucleotide Archive (https://www.ebi.ac.uk/ena), accession SRR14741640, were included in the clustering analysis.

All sequences were filtered by quality using a cut-off value of 20 and 90% of bases equal or above this value using the FASTQ/A short-reads pre-processing tools (Gordon 2010; http://hannonlab.cshl.edu/fastx_toolkit/) implemented in RepeatExplorer. The similarity-based clustering analysis was performed using the RepeatExplorer pipeline provided by the ELIXIR-CZ project part of the international ELIXIR infrastructure (https://www.repeatexplorer-elixir.cerit-sc.cz) (Novák et al. 2010, 2013, 2019).

An individual analysis to better characterize all TE families (Table 1), was performed in RepeatExplorer2. This analysis followed the established protocol of Novák et al. (2020) and employed default settings. The sequences were sampled to reach a genomic coverage of 0.1x selecting 2.5M reads for each sample during the automatic sampling step, so that the number of reads analyzed was proportional to the genome sizes (Table 1). The final coverage used for clustering after automatic sampling by RepeatExplorer was calculated as follows: coverage = (r×l)/g, where r corresponds to the number of analyzed reads after clustering, l to read length, and g to haploid genome size in bp.

**Table 1.** Genome sizes, number of automatically sampled reads for the clustering analyses, and resulting coverage for the analyzed tree tomato populations.

|  | **286** | **899** | **900** | **901** | **903** | **DB** |
| --- | --- | --- | --- | --- | --- | --- |
| **1C (Mpb)** | 11168,76 | 11168,76 | 11286,12 | 11212,77 | 11310,57 | - |
| **Reads long** | 150 | 150 | 150 | 150 | 150 | 150 |
| **Number of reads in the library** | 6,691,482 | 5,909,200 | 6,529,926 | 6,559,422 | 19,934,450 | 26,312,574 |
| **Individual clustering reads** | 1214367 | 1225987 | 1267472 | 1190448 | 1273672 | 2697492 |
| Coverage | 2,17 | 2,17 | 2,16 | 2,22 | 2,10 | - |
| Proportion de reads in top clusters | 61% | 60% | 60% | 61% | 60% | 66% |
| Singlets | 316570 | 328939 | 339023 | 313862 | 336532 | 575409 |
| Clusters | 48396 | 49376 | 50273 | 45858 | 49511 | 95183 |
| Superclusters | 48327 | 49307 | 50209 | 45789 | 49435 | 95112 |
| **Comparative clustering reads** | 852680 | 853626 | 852850 | 850708 | 853928 | 853598 |
| Total pb | 11168760000 | 11168760000 | 11286120000 | 11212770000 | 11310570000 | - |
| Total de reads | 74458400 | 74458400 | 75240800 | 74751800 | 75403800 | - |
| Coverage | 0,29 | 0,29 | 0,28 | 0,29 | 0,28 | - |

TEs were automatically annotated using the green plants (Viridiplantae) database implemented in RepeatExplorer2. A database of retrotransposon protein domains (REXdb) is implemented in the RepeatExplorer web server which provides a classification system that reflects the phylogenetic relationships and distinct sequence and structural features of the elements (Neumann et al. 2019). Resulting clusters with genomic percentages above 0.01% were further examined manually to characterize the most abundant repetitive families. Unclassified clusters were analyzed using similarity searches with BLASTN and BLASTX against the non-redundant protein databases (https://www.blast.ncbi.nlm.nih.gov/Blast.cgi). Repeat composition was calculated excluding clusters corresponding to organellar DNA, which represents extranuclear DNA from chloroplasts and mitochondria.

#### Comparative analysis of repeats

A comparative repeat analysis was performed in RepeatExplorer2, following the protocol described by Novák et al. (2020) and employing default settings. This approach involved the simultaneous clustering of sequencing reads from all six *S. betaceum* populations to identify shared elements and assess differences their relative abundances. Interlace reads from each accession were identified with a different prefix, sampled for 0.1× genome coverage each, concatenated and clustered in the same pipeline using the comparative analysis option (Table 1). The distribution of the 225 most abundant comparative repeat clusters, was graphically represented, excluding clusters representing plastid-derived sequences.

### Satellite Analysis

The same dataset used in the clustering analyses was run with the TAREAN (Tandem Repeat Analyzer) tool, implemented in RepeatExplorer2 server. TAREAN identifies putative tandemly arranged repetitive DNAs and reconstruct consensus sequences of putative satDNAs based on k-mer analysis (Novák et al. 2017). The similarity and identity of satellite monomers accross different samples were compared via local alignment (Table 2), using the MAFFT algorithm implemented in Geneious Prime (Kearse, 2012). Tandem organization was further inspected using dot plot analyses with default parameters and GC percentage of consensus monomers was determined, both within Geneious Prime (Kearse, 2012). Satellites were classified according to Ruiz-Ruano et al. (2016). Satellites with >95% identity were considered the same variant; those with 80-95% identity were considered different variants within the same family; and those with <80% identity were classified as different families.

**Table 2.**
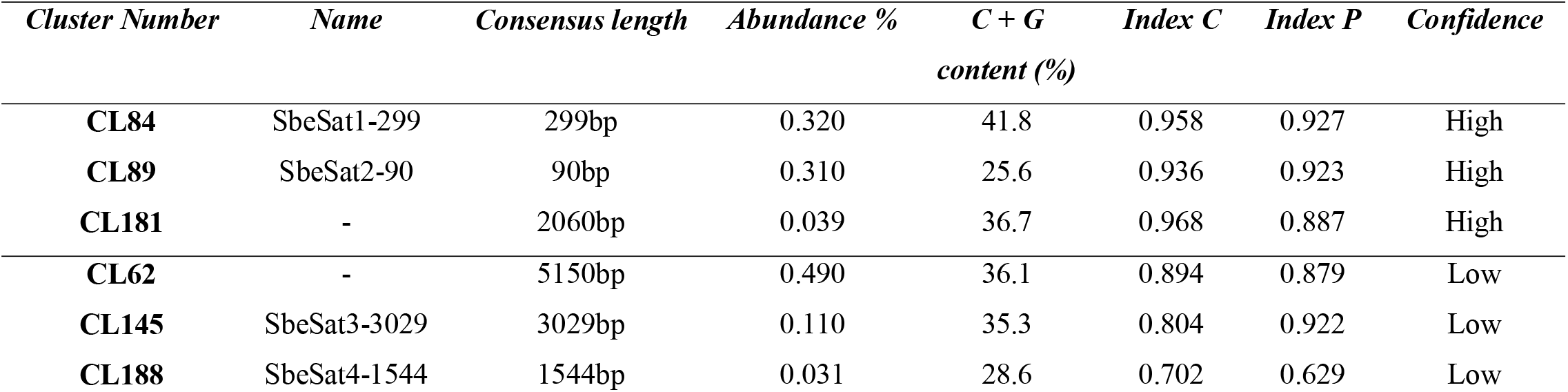
Main satellite DNAs identified by TAREAN.

### Cytogenetic Techniques

Mitotic chromosome preparations were obtained from root meristems pretreated with 2 mM 8-hydroxyquinoline for 6 h at 14°C and fixed in ethanol/acetic acid (3:1, v:v). Tissues were digested with Pectinex enzyme solution (Novozimes) and squashed in 45% acetic acid. Preparations were frozen in liquid nitrogen to remove the coverslips. For CMA/DAPI staining, slides were aged for 3 days, stained with CMA 0.1 mg/mL for 30 min, and mounted in glycerol/McIlvaine buffer pH 7.0 (1:1) containing 2.5 mM MgCl2 and 1 μg/mL DAPI. Slides were maintained in the dark at room temperature for 3 days before analysis.

Fluorescence *in situ* hybridization (FISH) was carried out to detect satellite DNA patterns, following protocols by Schwarzacher and Heslop-Harrison (2000). To reach stringency above 76%, the hybridization mix had 2x SSC, 50% v/v Formamide, 20% v/v dextran sulfate, 0.1% v/v SDS, and 4–6 ng/μL probes. Post-hybridization washes consisted of 2x SSC, 0.1x SSC, and 2× SSC, for 10 min at 42 °C each. Probes were obtained by PCR as described in González et al. (2020), the integrity of the PCR products was verified by electrophoresis in an agarose gel and after labeled with biotin (Bionick, Invitrogen) or digoxigenin (DIG Nick translation mix, Roche). The detection was made with Avidin-FITC (Sigma-Aldrich) and antiDIG antibodies conjugated with rhodamine (Roche). Chromosome metaphases were photographed using an Olympus BX61 microscope coupled with a monochromatic camera and Cytovision software (Leica Biosystems), and the images were pseudo-colored. The 5S and 35S rDNA probes used were designed by Waminal (2018) and synthetized and labeled in Macrogen (SouthKorea). The number and position of satDNA sites were determined by analyzing a minimum of ten selected metaphases across at least three slides per accession.

## Results

Flow cytometry analysis revealed that the estimated genome sizes of the different *Solanum betaceum* populations ranged from 1C 11.42 to 11.56pg (Fig1) representing a difference of approximately 140 Mbp.

To better characterize the repetitive fraction of the genome, a total of 5,138,161 random of 100 bp reads were analyzed using RepeatExplorer2, and representing ∼ 0.02 × coverage of the nuclear genome, after the removal of cytoplasmic DNA. Of these, 685,369 reads were classified as singlets, while 4,452,792 reads were grouped into 123,293 superclusters and 123,391 clusters. Among these, 258 clusters showed an abundance greater than 0.01% and were further analyzed. Overall, the repetitive fraction was estimated to account for 63.5% of the *S. betaceum* genome (Fig2).

**Figure 2.**
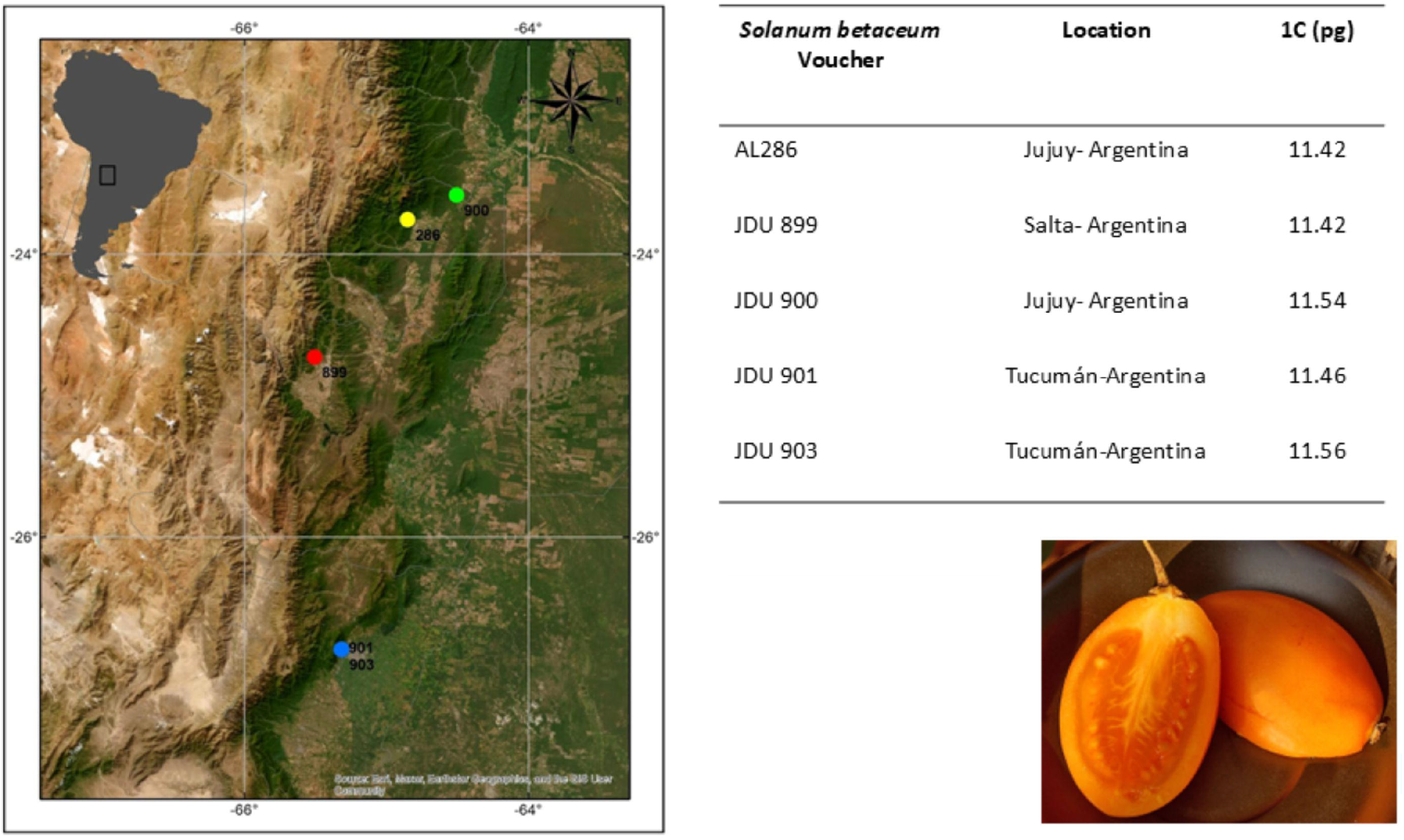
Genomic proportions (%) of repetitive sequences identified in *Solanum betaceum* after individual analysis in RepeatExplorer2

### Transposable elements

The annotated proportions of repetitive DNA of the analyzed samples are shown in Fig2. Retrotransposons were the most abundant components, representing □ 60% of the genome. Among these, Ty3-gypsy were particularly prevalent including the lineages Chromo, Ogre, and Athila groups, with the Tekay element being the most abundant, accounting for 44.3% of the genome. Ty1-copia elements represented 3.22% of the genome and included the lineafes Angela, Bianca, Ikeros, Ivana, SIRE, TAR, and the most abundant Tork (0.46%) groups (Fig2).

### Tandem repeated elements: Satellite DNA and rDNA

TAREAN analysis allowed the detection of tandem repeats associated with Satellite DNA and rDNA, which represented only 0.93% and 0.30% of the genome, respectively. The analysis identified three tandem repeat clusters (CL84, 89, and 181) with high confidence, and three with low confidence (CL62, 145, and 188). Clusters 181 and 62 showed sequence similarity with Ty3/gypsy-Tekay and Ty1/copia-Tork respectively. Cluster 84 (designated SbeSat1-299) represented 0.32% of the genome, with a satellite probability of 81.5%, a had a consensus monomer length of 299 bp with a GC content of 41.8%. Cluster 89 (SbeSat2-90) represented 0.31% of the genome, with a satellite probability of 15%, a consensus monomer sequence of 90 bp in length, and a GC content of 21.6%. Cluster 145 (SbeSat3-3029) represented 0.11% of the genome, with a satellite probability of 7%, a consensus sequence of 3029 bp in length, and a GC content of 35.3%. Cluster 188 (SbeSat4-1544) represented 0.03% of the genome, with a satellite probability of 6%, a consensus sequence of 1544 bp in length, and 28.6% GC content. The rDNA fraction was estimated to represent 0.29% of the genome for 35S and 0.01% for 5S (Table 2; Fig3).

**Figure 3.**
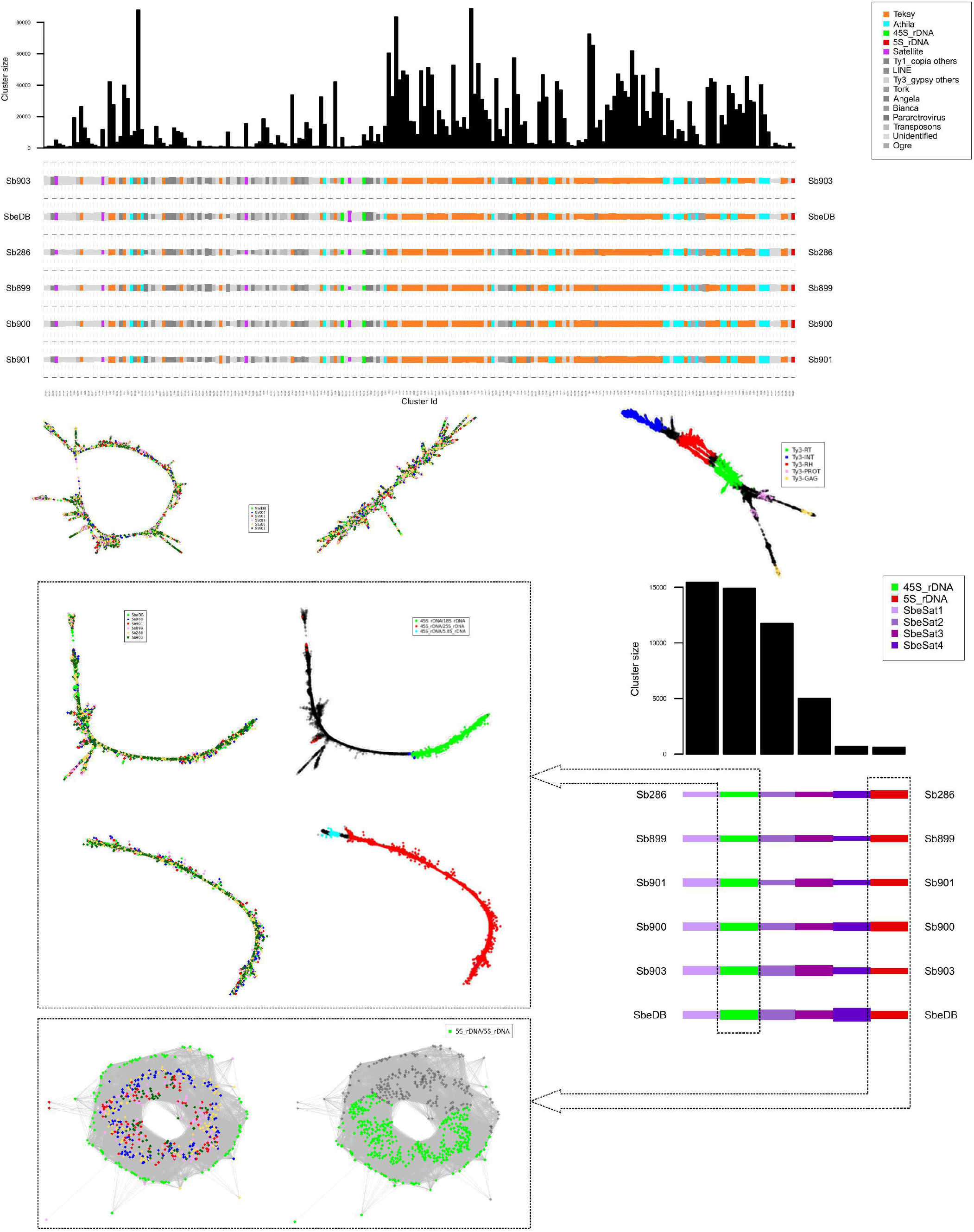
Genomic proportions (%) of repetitive sequences identified in different populations of *Solanum betaceum* through comparative analysis using RepeatExplorer2

#### Comparative analysis of repeats

To further characterize the repetitive fraction across different populations, we performed a comparative repetitive sequence analysis. The overall proportions of repetitive sequences observed were consistent with those obtained in the individual analyses. SCL1 was classified as Ty3/gypsy-Tekay. The greatest difference was found in the Tekay fractions, which range from 46.83% to 51.36% and in the satellite DNA fraction, which varied from 0.904% to 1.66% among populations (Table 3; Fig3). Among the different satellites, SbeSat1-299 ranged from 0.21% (in Sb286) to 0.36% (in Sb903); SbeSat2-90 varied from 0.16% in Sb286 to 0.32% in Sb903; SbeSat3-3029 from 0.07% in Sb286 to 0.14% in Sb903 and SbeSat4-1544 from 0.007% (in Sb899) to 0.025% (in SbDB) (Fig3). The proportion of 5S rDNA was similar between all populations (0.01%), while the 35S rDNA showed more variability, ranging in abundances from 0.22% to 0.37% (Table 3).

**Table 3.** Genomic proportions (%) of repetitive sequences identified in *Solanum betaceum* after comparative analysis in RepeatExplorer2.

| Comparative analysis | Sbe286 | Sbe899 | Sbe900 | Sbe901 | Sbe903 | SbeDB |
| --- | --- | --- | --- | --- | --- | --- |
| **Class I** | 0,20 | 0,24 | 0,23 | 0,21 | 0,32 | 0,29 |
| pararetrovirus | 0,35 | 0,37 | 0,37 | 0,39 | 0,39 | 0,50 |
| LINE | 0,61 | 0,68 | 0,56 | 0,66 | 0,62 | 0,55 |
| **LTR** | 2,41 | 2,42 | 2,51 | 2,44 | 2,47 | 2,41 |
| **LTR/Ty1_copia** | 0,01 | 0,01 | 0,01 | 0,01 | 0,01 | 0,01 |
| Ale | 0,03 | 0,04 | 0,03 | 0,03 | 0,03 | 0,03 |
| Alesia | 0,04 | 0,04 | 0,04 | 0,04 | 0,04 | 0,04 |
| Angela | 0,44 | 0,44 | 0,43 | 0,43 | 0,51 | 0,49 |
| Bianca | 0,42 | 0,41 | 0,42 | 0,38 | 0,51 | 0,45 |
| Ikeros | 0,06 | 0,07 | 0,07 | 0,06 | 0,09 | 0,08 |
| Ivana | 0,12 | 0,11 | 0,12 | 0,12 | 0,14 | 0,13 |
| SIRE | 0,14 | 0,14 | 0,14 | 0,12 | 0,15 | 0,15 |
| TAR | 0,39 | 0,42 | 0,41 | 0,40 | 0,52 | 0,50 |
| Tork | 0,52 | 0,53 | 0,52 | 0,49 | 0,53 | 0,54 |
| **Total Ty1_copia** | **2,17** | **2,21** | **2,19** | **2,08** | **2,53** | **2,42** |
| **LTR/Ty3_gypsy** |  |  |  |  |  |  |
| Chromovirus | 0,36 | 0,33 | 0,38 | 0,36 | 0,37 | 0,35 |
| Chromovirus/CRM | 0,11 | 0,12 | 0,13 | 0,10 | 0,14 | 0,14 |
| Chromovirus/Galadriel | 0,06 | 0,05 | 0,07 | 0,05 | 0,05 | 0,06 |
| Chromovirus/Tekay | **50,57** | **50,02** | **50,85** | **51,36** | **46,83** | **48,27** |
| Non-chromovirus/OTA/Athila | **7,85** | **7,67** | **7,35** | **8,24** | **7,16** | **7,13** |
| Non-chromovirus/OTA/Tat/Ogre | 0,98 | 0,98 | 0,92 | 0,91 | 0,97 | 1,02 |
| **Total Ty3_gypsy** | **59,93** | **59,17** | **59,7** | **61,02** | **55,52** | **56,97** |
| **Class II** |  |  |  |  |  |  |
| Subclass_1/TIR/EnSpm_CACTA | 0,11 | 0,11 | 0,12 | 0,11 | 0,15 | 0,13 |
| Subclass_1/TIR/hAT | 0,14 | 0,14 | 0,14 | 0,14 | 0,18 | 0,19 |
| Subclass_2/Helitron | 0,04 | 0,04 | 0,05 | 0,04 | 0,06 | 0,06 |
| **Total Class II** | **0,29** | **0,29** | **0,31** | **0,29** | **0,39** | **0,38** |
| **rDNA** |  |  |  |  |  |  |
| 45S rDNA | 0,22 | 0,25 | 0,30 | 0,31 | 0,30 | 0,37 |
| 5S rDNA | 0,01 | 0,01 | 0,01 | 0,01 | 0,01 | 0,01 |
| **Satellite** | 0,904 | 1,12 | 1,27 | 1,22 | 1,66 | 1,54 |
| **Total Tandem repeats** | **1,134** | **1,38** | **1,58** | **1,54** | **1,97** | **1,92** |
| **Unclassified** | 3,74 | 3,80 | 3,85 | 3,82 | 4,36 | 4,25 |
| **TOTAL** | **70,46** | **70,08** | **70,70** | **71,95** | **67,81** | **68,97** |

#### Chromosomal distribution of repetitive DNAs

To determine the distribution of heterochromatin (CG-rich regions), we performed double staining with the fluorochromes CMA and DAPI. No complex heterochromatin pattern or DAPI+ bands were observed with this procedure. However, we identified three chromosome pairs with terminal CMA+ bands (Fig5). Regarding the rDNA sites, *S. betaceum* showed one pair of interstitial 5S rDNA site and two pairs of terminal 35S rDNA sites located on the short arms of a pair of metacentric chromosomes, and co-localized with two of the three terminal CMA+ band (Fig.4).

**Figure 4.**
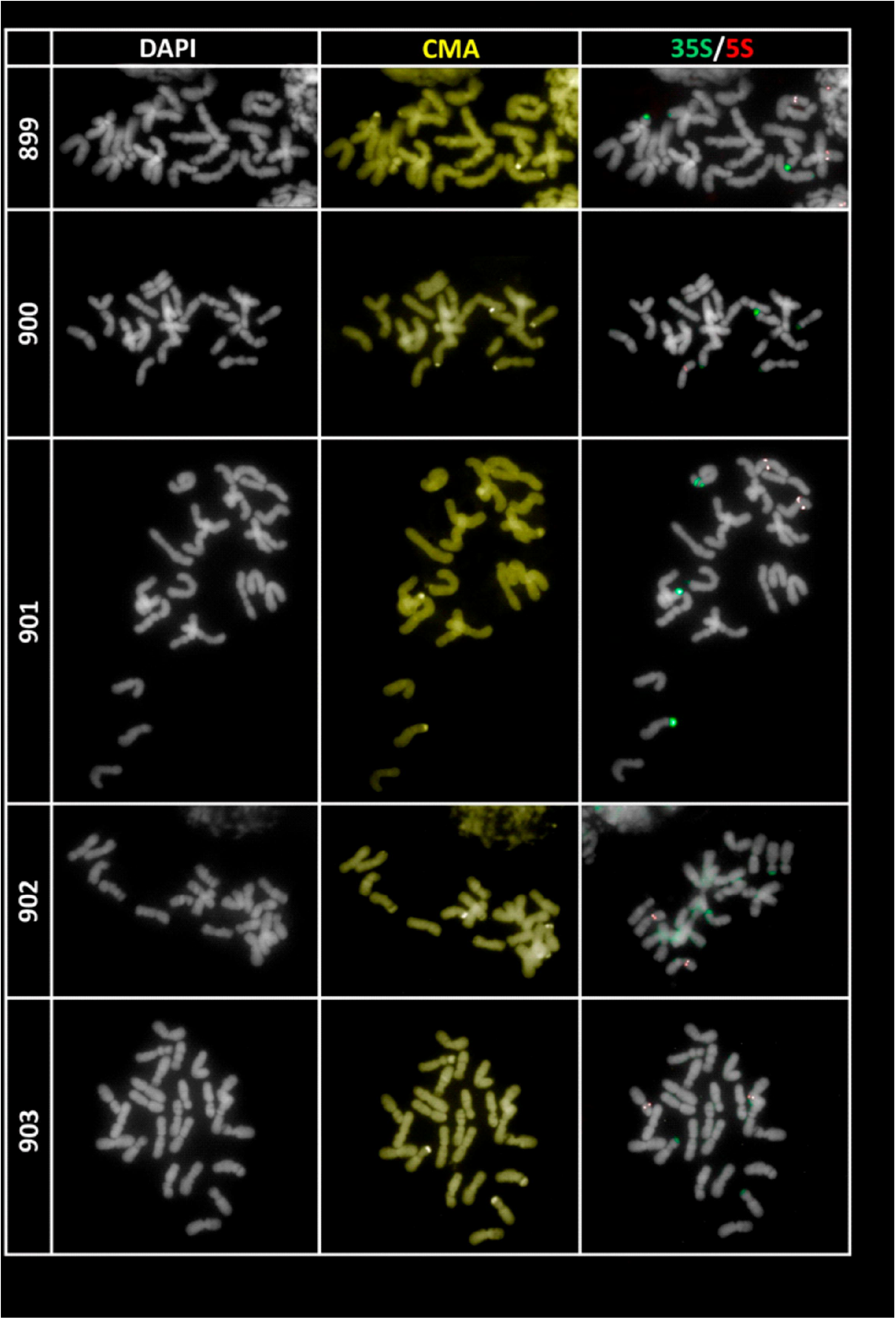
Mitotic metaphase chromosome spread displaying the CMA banding pattern and ribosomal DNA distribution

**Figure 5.**
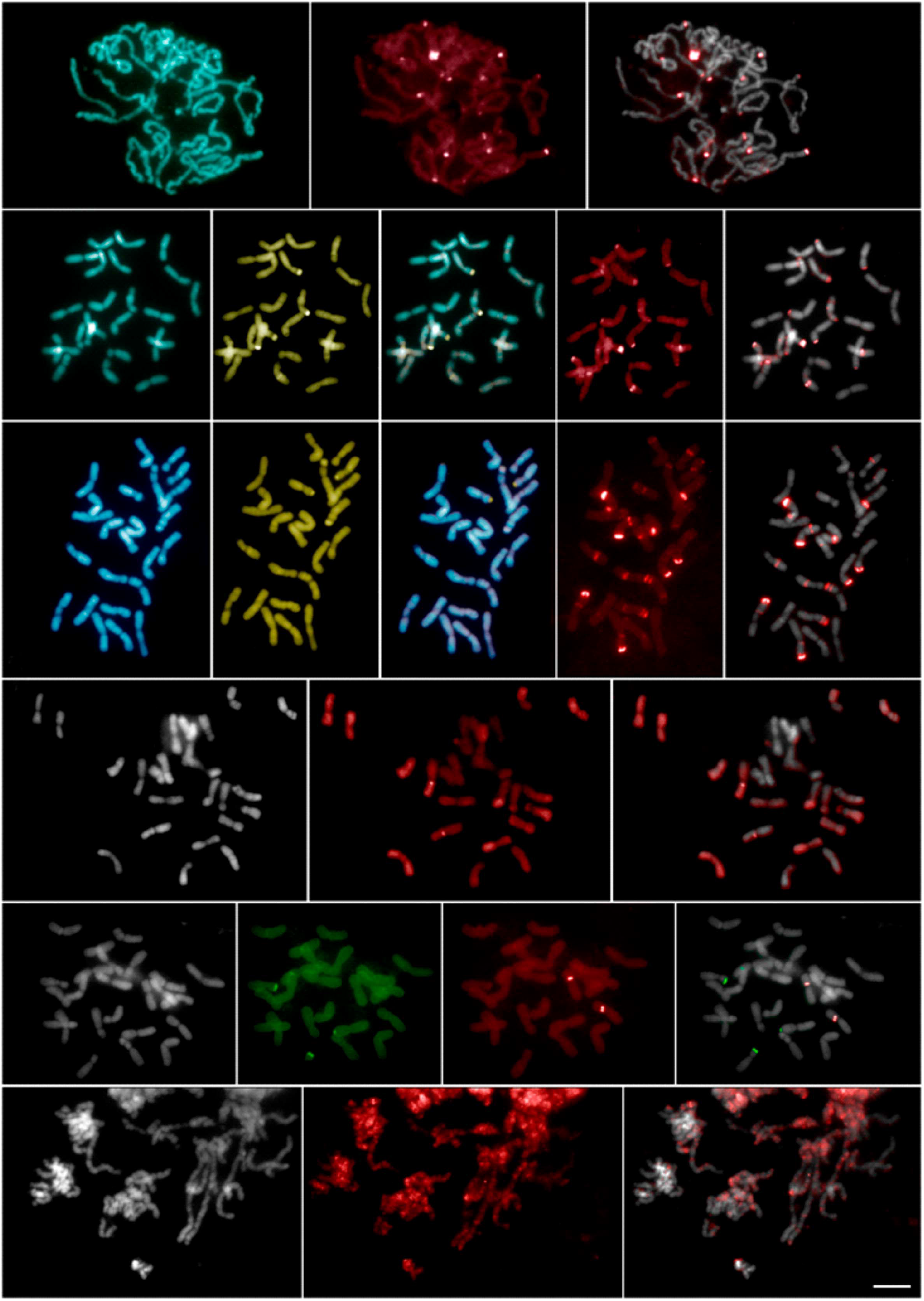
Mitotic metaphase chromosome spread displaying the SatDNA distribution

FISH analysis revealed that the satellite SbeSat1-299 showed signals in terminal and interstitial regions of eight chromosome pairs, co-localized with one pair of terminals CMA+ band that did not correspond to the 35S rDNA (Fig. XX). The satellite DNA SbeSat2-90 was detected on ten chromosome pairs in terminal, interstitial, and pericentromeric regions and co-localized with SbeSat1-299 at one pair of terminal CMA+ bands (Fig5). The satellite DNA SbeSat3-3029 showed one conspicuous interstitial block along with dispersed signals on several chromosomes (Fig5). Finally, SbeSat4-1544 showed a dispersed hybridization pattern across the chromosomes (Fig5). All four satellite families showed hybridization signals in all populations.

## Discussion

### Genome Size and Repeatome in Solanum betaceum

The Cyphomandra group (Bohs 2005, 2007; Weese and Bohs 2007; Särkinen et al. 2013; Gagnon et al. 2022), one of the 12 major evolutionary lineages within the highly diverse genus *Solanum* (∼1300 species, Gagnon et al. 2022), comprises approximately 50 species of woody shrubs or small trees distributed throughout the Neotropics. This lineage includes members of *Solanum* section *Pachyphylla* (Dunal) Dunal—previously treated as the distinct genus *Cyphomandra* Mart. ex Sendtn. (Bohs 1994, 1995)—as well as species of *Solanum* section *Cyphomandropsis* Bitter, as defined by Bohs (2001). Despite its taxonomic and ecological relevance, cytogenetic data are currently available for only around 40% of the species in this group, and genome size estimates exist for merely 24% (Chiarini et al. 2018).

One of the most striking features of the Cyphomandra clade is its notable divergence from most other *Solanum* species in both chromosome and genome size. In the majority of *Solanum* (excluding the Cyphomandra group), chromosomes are relatively small—typically ranging from 1 to 3 µm (Bernardello and Anderson 1990; Bernardello et al. 1994; Acosta et al. 2005, 2012; Chiarini and Bernardello 2006; Rego et al. 2009; Mesquita et al. 2019)—and genome sizes are modest, averaging 1C = 1.40 pg, with a range of 0.6 to 3.7 pg (Leitch et al. 2021). By contrast, species within the Cyphomandra clade possess significantly larger chromosomes, measuring between 4 and 10 µm (Pringle and Murray 1993; Acosta et al. 2012; Miguel et al. 2012; Moyetta et al. 2013; Mesquita et al. 2024), and exhibit substantially larger genomes, with an average 1C value of 10.71 pg and a range from 6.8 to 24.8 pg (Leitch et al. 2021).

To better understand the genomic architecture underlying these characteristics, we conducted the first *in silico* analysis of repetitive DNA in *Solanum betaceum*, a fruit crop of increasing agronomic interest within this clade. Our results revealed a genome size of 1C = 11.56 pg, which is consistent with previous estimates (Mesquita et al. 2024), reinforcing the distinctiveness of the Cyphomandra group in terms of nuclear DNA content. In our analysis, repetitive elements were found to constitute a substantial portion of the *S. betaceum* genome (63.5%), a percentage consistent with other angiosperm species characterized by large genomes, such as *Vicia faba, Allium cepa*, and *Bomarea edulis* (Macas et al. 2015; Peška et al. 2019; Nascimento et al. 2025). As expected, the repetitive landscape of *S. betaceum* is dominated by Ty3-gypsy long terminal repeat (LTR) retrotransposons (∼52%), with the Tekay lineage alone accounting for more than 40% of the repetitive fraction, followed by Athila elements (∼7%) and satellite DNA (∼1%). All other repeat families were found at lower abundance (≤1%).

Interestingly, although copia-type elements are less abundant in the genome (∼5%), they displayed higher variability, with nine distinct copia lineages identified. In contrast, six lineages were detected for Ty3-gypsy, indicating a somewhat lower diversification despite their numerical dominance. This pattern was also observed in other *Solanum* species such as potato and tomato (Gaiero et al. 2019), and may suggest that copia elements have undergone more recent insertions or are subject to more efficient elimination mechanisms, contributing to their greater lineage diversity.

The prominence of Ty3-gypsy retrotransposons, especially those belonging to the Tekay lineage, suggests that these elements have played a central role in the expansion and structural evolution of the *S. betaceum* genome. Similar patterns in other plant lineages—such as *Allium* (Peška et al. 2019), *Caesalpinia* (Van-Lume et al. 2019), and *Passiflora* (Sader et al. 2021)—underscore the broader significance of specific retroelement lineages in driving genome size dynamics across angiosperms. Moreover, our results align with findings from *Solanum* species by Gaiero et al. (2019), where the amplification of retrotransposons, particularly Ty3-gypsy-Tekay, has been identified as a major driver of genome size variation.

The observed pattern of repetitive DNA in *S. betaceum* contrasts with that of species with smaller genomes, where a higher proportion of satellite DNA is often present. This difference may reflect a contrasting strategy in genome organization, where in smaller genomes, satellite DNA plays a larger role in forming heterochromatin blocks, contributing to a compact chromosomal architecture. In contrast, species with larger genomes, such as *S. betaceum*, appear to rely more on the expansion of retrotransposons, which may lead to the dispersion of heterochromatin across the genome, contributing to overall genome expansion. Taken together, these findings strongly support the hypothesis that Tekay and Athila elements are major contributors to the evolution of the large genome observed in *S. betaceum*, highlighting the importance of these elements in the evolution of the genomes of several species of the family.

### Satellite DNA Composition and Cytogenetic Potential

In contrast to the high abundance of retrotransposons, satellite DNA and ribosomal DNA (rDNA) were found in much lower proportions, representing 0.93% and 0.30% of the genome, respectively. This pattern aligns with general trends observed in Solanaceae, where retroelements typically dominate over tandem repeats in terms of genomic representation (Garrido-Ramos, 2017; Gaiero et al., 2019). Nonetheless, the identification of four distinct satellite DNA families (SbeSat1-299, SbeSat2-90, SbeSat3-3029, and SbeSat4-1544) enhances our understanding of chromosomal organization and provides promising cytogenetic markers. A low diversity of satellite DNA families has also been reported in other species with large genomes. For instance, in *Passiflora*, satellite DNA diversity decreases with increasing genome size (Sader et al., 2021).

In our double-staining analyses using CMA and DAPI fluorochromes, we detected three pairs of terminal CMA□ bands, two of which co-localized with 35S rDNA sites, and a third that co-localized with SbeSat1-299 and SbeSat2-90. These findings indicate that heterochromatic bands in *S. betaceum* are composed of 35S rDNA and two distinct satellite DNA families. Among them, SbeSat1-299, the most abundant, has a higher GC content (45%) compared to SbeSat2-90 (25%). Both satellite DNAs are distributed across terminal, interstitial, and pericentromeric regions, forming conspicuous bands even within euchromatic areas. Similar patterns have been reported in previous studies, where satellite DNAs are not restricted to heterochromatin but are also found in euchromatin (Rico-Porras et al., 2024; Nascimento et al., 2025). Interestingly, several satellite DNA sequences exhibited similarity to transposable elements, such as members of the Tork, CACTA, Retand, and Angela lineages. This suggests a possible origin from transposable elements, a phenomenon reported in other plant species as well (Belyayev et al., 2020), or tandem-repeats carrying Tes as suggested by some authors (Navarro-Dominguez et al., 2023). Both possibilities, reinforce the dynamic interplay between different classes of repetitive DNA in genome evolution (Šatović-Vukšić and Plohl, 2023).

The classical view of satellite DNAs involves their organization into clustered arrays that form heterochromatic blocks. However, as more satellite DNAs are identified and mapped onto chromosomes, a greater diversity in their distribution patterns has been observed. As in other plant genera (Ribeiro et al., 2022; Yücel et al., 2024), two satellite DNAs in *S. betaceum*, SbeSat3-3029 and SbeSat4-1544, exhibited a highly dispersed distribution throughout the chromosomes. Thus, although *S. betaceum* does not possess a large number of different satellites, it still displays the full range of satellite diversity in terms of genomic abundance, chromosomal distribution, association with transposable elements, and localization in both heterochromatin and euchromatin (Šatović-Vukšić and Plohl, 2023).

### Intraspecific Variation and Evolutionary Implications

Our analysis revealed intraspecific variation in genome size and the abundance of specific repetitive elements across different populations of *S. betaceum*. Although the total amount of repetitive elements did not correlate with differences in genome size, satellites and certain classes of retrotransposons followed the expected pattern. For instance, the sample with the higher DNA content exhibited nearly twice the abundance of satellite sequences. Notably, differences in Tekay and satellite DNA content point to a dynamic repeatome, shaped by processes such as geographical isolation, ecological adaptation, and domestication. This observation is in line with the idea that repetitive elements are not only structural genome components but also drivers of genetic and phenotypic diversity (Plohl et al. 2012; Feschotte and Pritham 2007).

The relatively low proportion of satellite DNA, compared to other species with large genomes (e.g., *Passiflora quadrangularis*), may reflect differences in genome architecture, regulatory constraints, or turnover rates of tandem repeats. Furthermore, the potential dispersion of satellite sequences in euchromatic regions—suggested in recent literature (Šatović-Vukšić and Plohl 2023)— may also apply to *S. betaceum* and deserves further cytological investigation.

The repetitive DNA landscape characterized here provides a foundation for the development of molecular tools for breeding and conservation. The satellite DNA sequences identified can serve as cytogenetic markers, aiding in karyotype characterization, genome mapping, and evolutionary studies. Moreover, the observed genomic diversity among populations supports the implementation of conservation strategies to preserve the genetic resources of this species, especially given its increasing importance as a fruit crop.

### Conclusions

Our study represents the first comprehensive analysis of the repeatome in *Solanum betaceum*. The genome is predominantly composed of Ty3-gypsy retrotransposons, with the Tekay lineage playing a central role. Although satellite DNA accounts for a minor proportion of the genome, it displays notable variation among populations, highlighting its potential as a cytogenetic marker and its involvement in genome evolution. These findings offer valuable insights into genome organization, contribute to the understanding of repetitive DNA dynamics in the Cyphomandra clade, and lay the groundwork for future breeding and conservation efforts in *S. betaceum*.

## Acknowledgments

We are grateful to the Consejo Nacional de Investigaciones Científicas y Técnicas (CONICET), the Agencia Nacional de Promoción Científica y Tecnológica (FONCyT), and SECyT (Universidad Nacional de Córdoba, Argentina) for their financial support, as well as for providing equipment and facilities. We also thank Dr. Andreas Houben for funding the Illumina population-level sequencing. We also thank the ELIXIR-CZ Research Infrastructure Project (LM2015047) for providing computational resources for RepeatExplorer analysis.

## Author Contributions

MAS designed experiments, performed bioinformatic analysis, amplifications and probe labeling, FISH, and data analysis, and wrote the manuscript. MV performed flow cytometry analysis, analyzed data, and assisted with writing the manuscript; FC, MJZ, and AL provided plant material and taxonomic identification. JDU designed the experiments, analyzed data, and wrote the manuscript. All authors have read and agreed to the published version of the manuscript.

